# A survey of opsin localization, glycosylation, and light/chromophore influence on degeneration in 26 *rhodopsin*-associated retinitis pigmentosa models

**DOI:** 10.1101/2025.07.23.665973

**Authors:** Aaron D. Loewen, Beatrice M. Tam, Colette N. Chiu, Ross T. Scharbach, Orson L. Moritz

**Affiliations:** Department of Ophthalmology and Visual Sciences, University of British Columbia, Vancouver, British Columbia, Canada. V5Z 3N9

## Abstract

**Purpose:** Mutations in *rhodopsin (RHO)* cause autosomal dominant retinitis pigmentosa (RP), which has multiple clinical subclasses, including class B1 (“sector”) RP in which the asymmetric retinal degeneration (RD) suggests an environmental influence. The pathogenic mechanisms of most class B1 mutations are uncharacterized. We generated new animal models of *RHO*-associated RP to examine RHO expression, localization, glycosylation, and the influence of light and chromophore binding on RD.

**Methods:** We generated transgenic *X. laevis* expressing wildtype or mutant human *RHO* transgenes. Confocal images were used to evaluate RD and trafficking. Immunoassays were used to quantify RD and investigate RHO glycosylation.

**Results:** We created *X. laevis* models of 26 different forms of *RHO*-associated RP. Most mutations caused RD, with the exception of those at residue R135 and G101. Many variants did not alter RHO localization. Multiple class B1-associated RHO mutants induced light-dependent RD, suggesting light is the environmental influence associated with the class B1 phenotype. However, the degeneration associated with two partially ER-retained class B1 mutants (S22R and D190G) was not mitigated by dark rearing. P23H and S176F constituted a distinct subclass associated with inner segment retention and proteolytic cleavage.

**Conclusions:** Many *RHO* mutations do not substantially alter RHO localization or glycosylation. The exceptions we identified are P23H and S176F, which dramatically mislocalize, and constitute a distinct category of proteolytically-cleaved misfolding variants. L31Q and T58R induce RD by mechanisms similar to glycosylation-deficient variants, despite lack of glycosylation defects. Intermediate phenotypes indicate at least one previously undescribed mechanism for class B1 RP pathogenesis.

## Introduction

Retinitis pigmentosa (RP) is an inherited retinal disease involving progressive blindness due to rod loss and subsequent cone death.^1^ There is no cure for RP. Vitamin A has been suggested as a treatment option, but has limited value in most cases.^2,3^ Other therapeutic strategies under investigation include pharmacological chaperones, increasing BiP expression, and gene editing.^4–8^

Mutations in *rhodopsin* (*RHO*) are responsible for one third of autosomal dominant (ad) RP cases.^9^ Over 200 *RHO* mutations are associated with adRP (www.hgmd.cf.ac.uk; Accessed December 26, 2024). *RHO-*associated adRP has multiple clinical subclasses defined by the rate and pattern of disease progression.^10^ This includes class B1 (“sector”) RP, where disease progression is slower and less severe than class A (“classic”) RP, and restricted to maximum two quadrants of the retina, generally the naso-inferior quadrants.^7,10^ This asymmetric retinal degeneration (RD) strongly implies an environmental influence, as all rods express the same *RHO* gene. A likely candidate is light, with evidence including light-dependant RD in transgenic animals with class B1 mutations, and disease patterns corresponding to light exposure histories.^11–15^ The phenotype implies that preventative treatment is possible.^16^

The most extensively studied class B1 *RHO* mutations are T4K, T17M, and P23H. P23H *RHO* causes protein misfolding and upregulation of the unfolded protein response, leading to cell death.^17–22^ P23H RHO prominently mislocalizes to the endoplasmic reticulum (ER) of rod inner segments (RIS) in animal models.^12,23^ In contrast, the localization of class B1-associated mutants T4K, N15S, and T17M RHO is indistinguishable from WT.^13,24^ RD is initiated upon photoactivation of T4K or T17M RHO in rod outer segments (ROS).^13,25^ These mutations comprise a mechanistic class distinct from P23H.^26^ Critically, therapeutic strategies for one class are likely to be ineffective or even detrimental for another.^25,27^

Previously established *RHO*-associated adRP classifications rely on cell culture or patient data.^10,28–30^ Many mutations have been classified in these systems but few have been examined in photoreceptors. Much research has focused on P23H, despite other mutations being more prevalent outside North America.^31–33^ It is unclear whether P23H disease mechanisms are representative of class B1 RP and *RHO* mutations in general. Previously, we successfully modeled *RHO*-associated RP in *Xenopus laevis.*^13,18,34^ Transgenic *X. laevis* are easily generated, RD occurs rapidly, and mutant RHO can be localized using antibodies selective for mammalian RHO.^25,35,36^ Therefore, to determine whether other *RHO* mutations cause misfolding in rods (similar to P23H), or cause photoactivation-induced cell death (similar to T4K and T17M), and whether additional mechanisms exist, we characterized 26 adRP-linked *RHO* mutations in transgenic *X. laevis* rods, including localization, glycosylation status, and influence of lighting conditions and chromophore binding on RD. Twelve of these mutations were class B1, ten were class A, and four were unclassified. We identified multiple mutations that cause photoactivation-induced cell death and demonstrate that the P23H phenotype of mislocalization and cleavage is shared by only one mutation, S176F. We also provide evidence of novel mechanisms.

## Methods

### Molecular biology

Transgene constructs were generated as previously described.^18^ Mutations were introduced into the human RHO cDNA, which was inserted into a XOP0.8-eGFP-N1 plasmid backbone, replacing the eGFP.^18,33^ Transgene constructs were linearized with FseI (NEB, USA). <u>Generation and rearing of transgenic *Xenopus laevis*</u>

Transgenic *X. laevis* tadpoles were generated as previously described.^18,36^. Healthy embryos of random sex were raised at 18°C on a 12-hour light cycle unless otherwise specified. Embryos were exposed to G418 (VWR, USA) after one day post-fertilization (dpf) for 4-6 days to eliminate non-transgenic embryos.^37^ Normally-developed *X. laevis* were sacrificed at 14 dpf (stage 48). One eye was fixed in 4% paraformaldehyde-3% sucrose, while the other was solubilized in SDS-PAGE loading buffer. F1 tadpoles were generated by mating WT males with transgenic females. F1 offspring were raised and sacrificed as described above but without G418 selection, in order to produce both transgenic and control non-transgenic siblings.

### Ethics approval

These procedures were approved by The University of British Columbia Animal Care Committee and conducted in accordance with the ARVO Statement for the Use of Animals in Ophthalmic and Vision Research.

### Dot blots

Dot blots of solubilized eye extracts were performed as previously described.^18^ Membranes were probed for human or *X. laevis* rod opsin using either mAb 1D4 (mammalian RHO; 1:5,000 dilution; UBC-UILO, Canada) or mAb B6-30N (transgenic and endogenous RHO; 1:10 dilution), followed by IR-dye800-conjugated goat anti-mouse antibody (IR-800) (1:10,000 dilution; LI-COR Biosciences, USA).^38,39^, and imaged using a LI-COR Odyssey system. Control standards were solubilized COS-7 cells expressing human rod opsin and 14 dpf WT tadpole eyes.

### Western blots

*X. laevis* eye extracts were electrophoresed through 10% or 12% SDS-PAGE gels, transferred to Immobilon P membranes (MilliporeSigma, USA), and probed with mAb 1D4 (1:5,000 dilution) or mAb A5-3-12 (1:10 dilution) and imaged as described for dot blots.^39^ For deglycosylation, extracts were incubated with 100 U of PNGaseF (NEB, USA) in 50 mM NaPO_4_ pH 7.5, and 1% NP-40 detergent at 37°C for one hour.

### Immunohistochemistry and confocal microscopy

Fixed eyes were cryosectioned at 12 µm and labelled with mAb 2B2 (1:10 dilution, anti-mammalian RHO) or mAb 1D4 (1:5,000 dilution, anti-mammalian RHO) followed by cyanine 3-conjugated (Cy3) anti-mouse antibody (1:750 dilution; Jackson Immunoresearch, USA), or anti-calnexin pAb (1:50 dilution; StressGen, Canada) followed by cyanine5 (Cy5) anti-rabbit secondary (1:750 dilution; Jackson Immunoresearch, USA).^35,38^ Sections were counter-labelled with Alexa-488-conjugated wheat germ agglutinin (WGA) (1:100 dilution; Life Technologies, USA) and Hoechst 33342 (1:1,000 dilution; Sigma-Aldrich, USA) and imaged with a ZEISS LSM800 confocal microscope equipped with a 40X/NA1.2 water-immersion objective (ZEISS, Germany).

### Colocalization of RHO and post-Golgi membranes in the RIS

Using ZEISS Zen 2 software, we compared the WGA and anti-RHO channels in 1024×1024 (pixel size: 0.039 µm^2^) images of RISs. We set a threshold by analyzing the nuclear region, where no WGA- or Cy3-positive signal was expected to be present and calculated a Pearson’s correlation coefficient (PCC) for the two channels using unadjusted raw 16-bit data images. PCC values were normalized using the Fisher transformation.^40^

### Data analysis and figure assembly

Relative expression levels of human RHO were calculated from the ratio of 1D4/B630N labeling on dot blots as previously described.^18,26^ For statistical analyses, a log transformation was applied to the transgenic rod opsin and total rod opsin levels. Data were analyzed using either ANCOVA or Student’s *t-*test using GraphPad Prism 10. For figures, we adjusted antibody labeling linearly, while counter-labels WGA and Hoechst were adjusted with altered “gamma” to better reveal cellular morphology. Final figures were assembled using Adobe Photoshop.

## Results

### Most RP-associated *RHO* mutants expressed in *X. laevis* cause RD

We generated 26 animal models of RP using primary transgenic *X. laevis* carrying human *RHO* transgenes, including 12 class B1 mutations (Fig. 1; Supplementary Table S1). Of the 26 mutations, 23 had not been previously modeled *in vivo.* We considered RD confirmed when at least three animals were evaluated by microscopy, and at least two had visible RD. Absence of RD was considered confirmed when at least five animals were evaluated by microscopy and none had apparent RD, and there was also no evidence of RD by dot blot (reduced rod opsin signal). We found RD caused by class A *RHO* mutations G89D, S176F, E181K, D190N, and A346P (Fig. 2A-C). RD caused by G89D and E181K *RHO* was also assessed by dot blot assay, in which we determined that increased expression of mutant human RHO was correlated with decreased total RHO (ANCOVA: G89D, p<0.0001; E181K, p=0.0005) (Fig. 2A). In contrast, as previously described, the human WT RHO transgene did not cause RD as assessed by confocal microscopy and dot blot (Fig. 2F, H).^13,18,25,26^ We did not see significant RD using either assay for the mutations F45L (ANCOVA: p = 0.1997), R135G (ANCOVA: p=0.8813), R135L (ANCOVA: p=0.9491), or R135W (ANCOVA: p=0.3186) (Fig. 2A, C).

**Figure 1.**
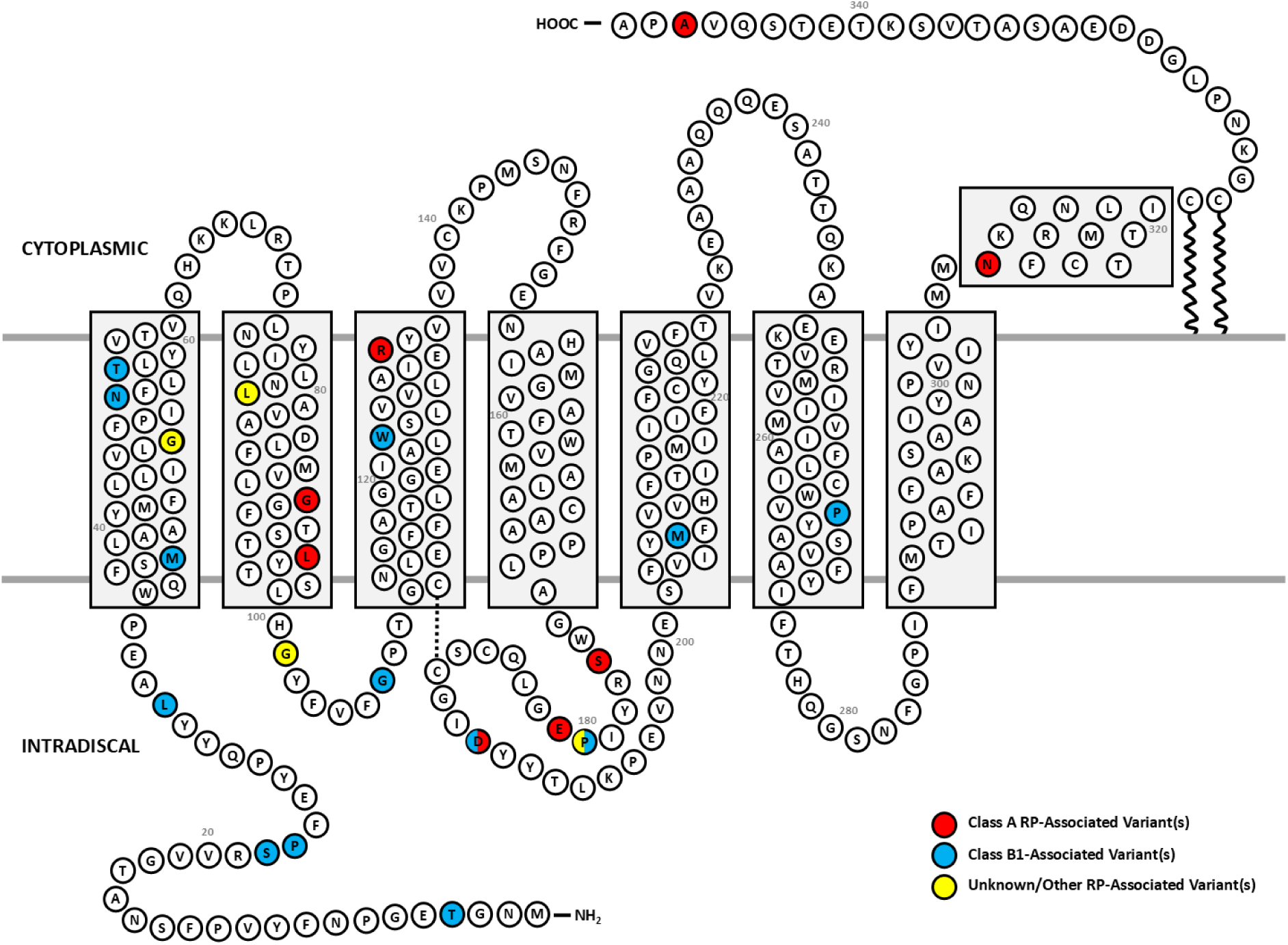
Locations of RP-associated *RHO* missense variants examined in this study. Coloured circles represent residues that, when mutated, have been reported to cause RP. Red indicates class A, blue indicates class B1, and yellow indicates RP mutations from an unknown or other class. Mutations listed in Supplementary Table S1.

**Figure 2.**
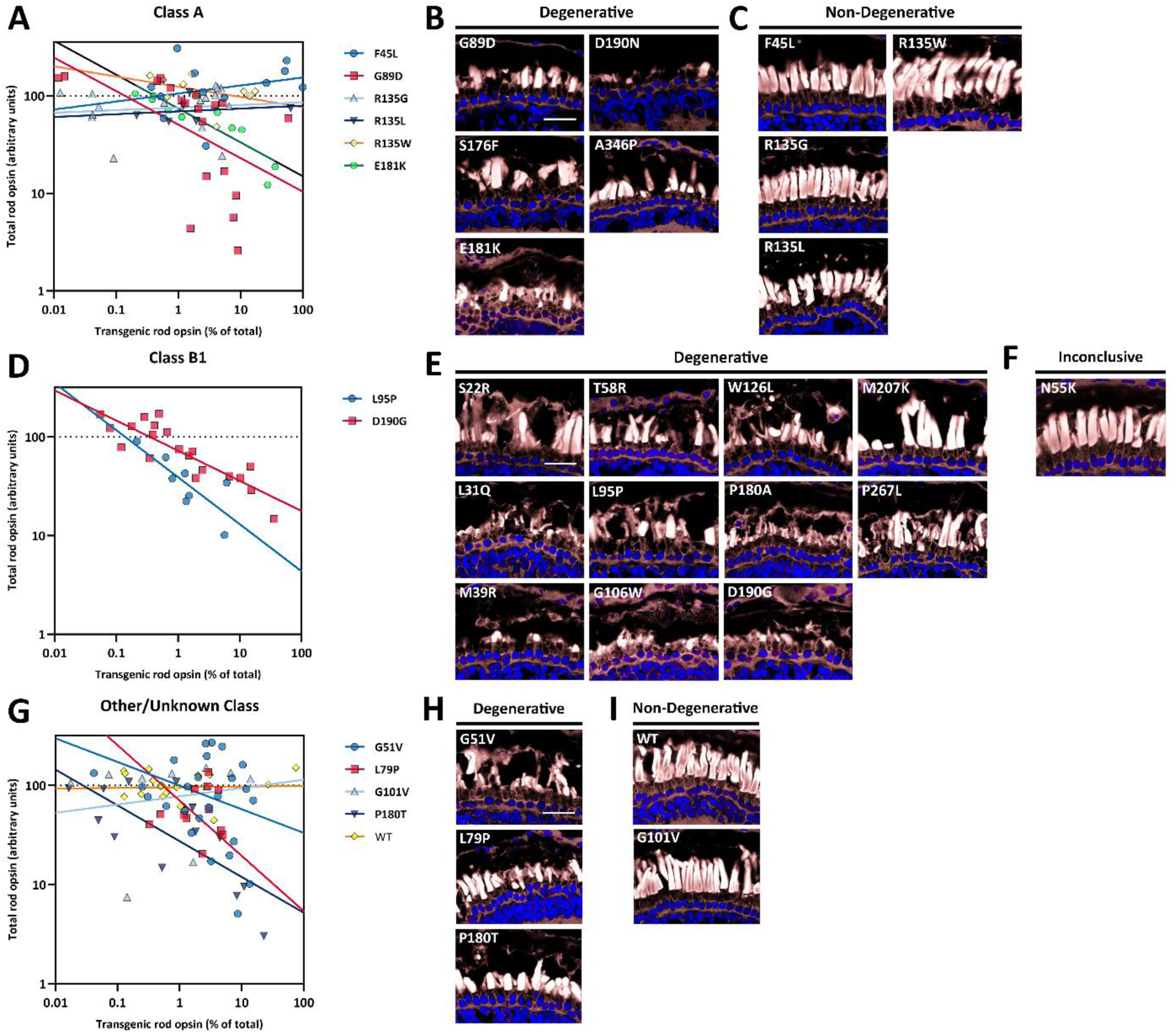
*X. laevis* expressing RP-associated RHO mutants show variable levels of RD. *A, D, G)* Dot blot results from solubilized retinas of 14 dpf *X. laevis* expressing different human *RHO* transgenes. Solid line indicates linear regression of log transformed dot blot data. N = 6-28 animals per genotype. *B, C, E, F, H, I)* Representative confocal images of transgenic *X. laevis* retinas showing RD (*B, E, H*), no RD (*C, I*), or inconclusive (*F*). White – AF488-WGA, blue – Hoechst. Bar = 25 μm.

For class B1 mutations, we found that S22R, L31Q, M39R, N55K, T58R, L95P, G106W, W126L, P180A, D190G, and M207K caused RD that was apparent by confocal microscopy (Fig. 2E). RD caused by L95P and D190G was also assessed by dot blot, and we determined that mutant RHO expression was correlated with decreased total RHO (ANCOVA: L95P, p=0.0129; D190G, p<0.0001) (Fig. 2D). We also generated animals expressing N55K RHO; however, the yield was insufficient to determine whether RD did or did not occur (Fig. 2F).

We also characterized four *RHO* mutations previously established as causative for RP, but with no assigned clinical class. We found that G51V, L79P, and P180T caused RD (ANCOVA: G51V, p=0.0073; L79P, p<0.0001; P180T, p<0.0001) (Fig. 2G-H). However, G101V did not cause RD by either assay (ANCOVA: p=0.8277) (Fig. 2G, I). Again, and as previously noted, human WT RHO did not cause RD at any expression level achieved by our system, indicating any RD observed is due to the presence of the *RHO* mutation (Fig. 2G, I).^12,13,18,24,25^ <u>RP-associated RHO mutants show a wide range of ER retention phenotypes</u>

We previously noted that transgenic *X. laevis* expressing the misfolding variant P23H RHO exhibited increased transgenic RHO labeling in the ER and decreased labeling in Golgi relative to those expressing human WT *RHO*.^12,18^ To quantify trafficking changes in RHO mutants, we assessed the presence of transgenic RHO in RIS and its colocalization with the Golgi as an indirect measure of ER-to-ROS trafficking. This method was used because of artifacts associated with anti-RHO labeling of ROS caused by high RHO concentrations.^12,41^ PCC values were generated from this colocalization measurement. Positive PCC values indicate ER-to-Golgi trafficking, while negative PCC values indicated biosynthetic defects and ER retention. Human WT and T4K RHO had average PCC values of 0.67 (95% CI: 0.5717 to 0.7748) and 0.57 (95% CI: 0.4805 to 0.6552) respectively (Fig. 3A). In contrast, human P23H RHO, which is retained in the ER, had an average PCC value of -0.1157 (95% CI: -0.1886 to 0.04292).

**Figure 3.**
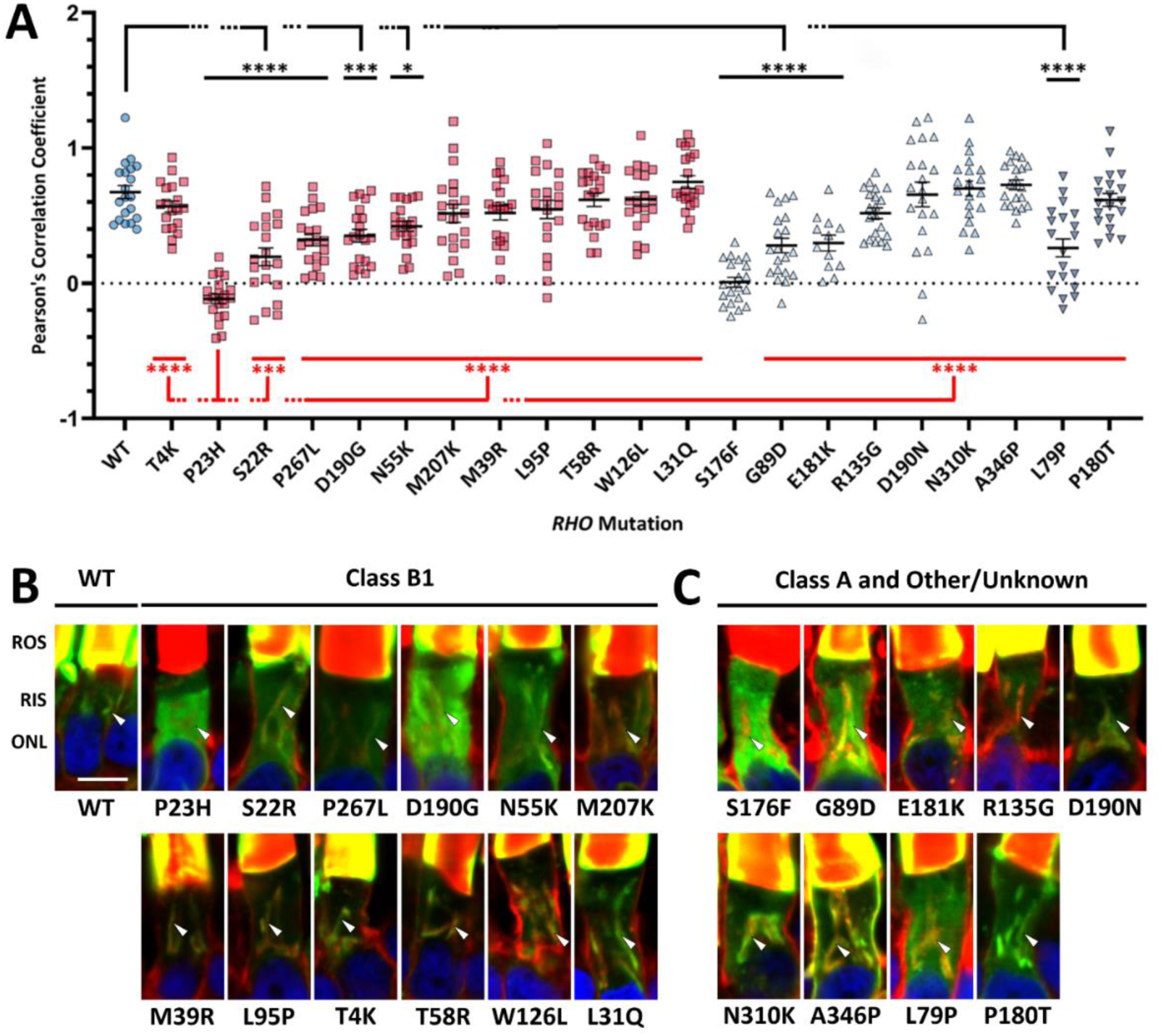
Colocalization analysis of RP-associated RHO variants with Golgi shows a variety of localization patterns. *A)* Pearson’s Correlation Coefficient (PCC) values after normalization correlating WGA and human RHO in the RIS (n = 12-20 photoreceptors taken from three to six animals per genotype). Circles – WT, squares – class B1, triangles – class A, inverted triangles – other/unknown class. Significance determined using the Dunnett’s multiple comparisons test to compare to human WT RHO (black bars) and P23H RHO (red bars) (see Supplementary Table S2 for p-values). Error bars = SEM. *B)* Confocal microscopy looking at rod inner segments and RHO localization of RP-associated RHO mutants. White arrows indicate the location of the Golgi body. Red – WGA, green – mAb 2B2 (anti-mammalian RHO), blue – Hoechst. ROS – rod outer segment, RIS – rod inner segment, ONL – outer nuclear layer. Bar = 5 μm.

We applied the imaging paradigm across RP-associated RHO mutants and compared results to WT and P23H. One-way ANOVA analysis determined colocalization values for *RHO* variants were non-uniform (p<0.0001). Post-hoc tests demonstrated that five class B1 variants (S22R, P23H, N55K, D190G, and P267L) had significantly lower colocalization with Golgi than WT (Fig. 3A-B). Seven class B1 mutants, including T4K, were not significantly different from WT. We also found varying degrees of ER retention for class A mutants, with three variants (G89D, S176F, and E181K) significantly different from WT, and four variants (R135G, D190N, N310K, and A346P) not significantly different from WT (Fig. 3A, C). Note that although A346P was prominently mislocalized to the plasma membrane, consistent with disruption of the C-terminal OS localization signal, this form of mislocalization was not captured by the ER retention measurements made in Figure 3.^34,41^ No other variants were mislocalized to the plasma membrane. Two variants of unknown class were investigated (L79P and P180T); only L79P had an altered localization (Fig. 3A). All mutants investigated except S176F had PCC values significantly higher than P23H (Fig. 3A). This was also readily apparent by visual inspection of images (Fig. 3B-C). Thus, P23H and S176F form a subclass in which trafficking is severely affected.

### Class B1 RP-associated *RHO* mutations outside of consensus glycosylation sites do not alter RHO glycosylation

The class B1 mutations T4K, N15S and T17M mutations alter the consensus N-linked glycosylation sites found at N2 or N15, and are associated with light-exacerbated RD.^13^ Thus, loss of glycosylation is a potential factor in the associated RD mechanism.^26,42^ In addition, biosynthetic defects such as those associated with P23H RHO can result in immature “high mannose” glycosylation states if the proteins are retained in the ER.^12^ To determine whether other class B1 mutations without trafficking defects alter RHO glycosylation, we deglycosylated retinal extracts with PNGase F and examined them by western blotting. A large shift in mobility between untreated RHO samples and deglycosylated samples indicates the presence of dual N-linked glycosylation. In contrast, natively mono-glycosylated mutants such as T4K have an altered mobility, but an identical mobility after deglycosylation. We examined four class B1 mutants (S22R, L31Q, T58R, and D190G) located outside consensus N-linked glycosylation sites and found that they did not alter RHO glycosylation (Fig. 4).

**Figure 4.**
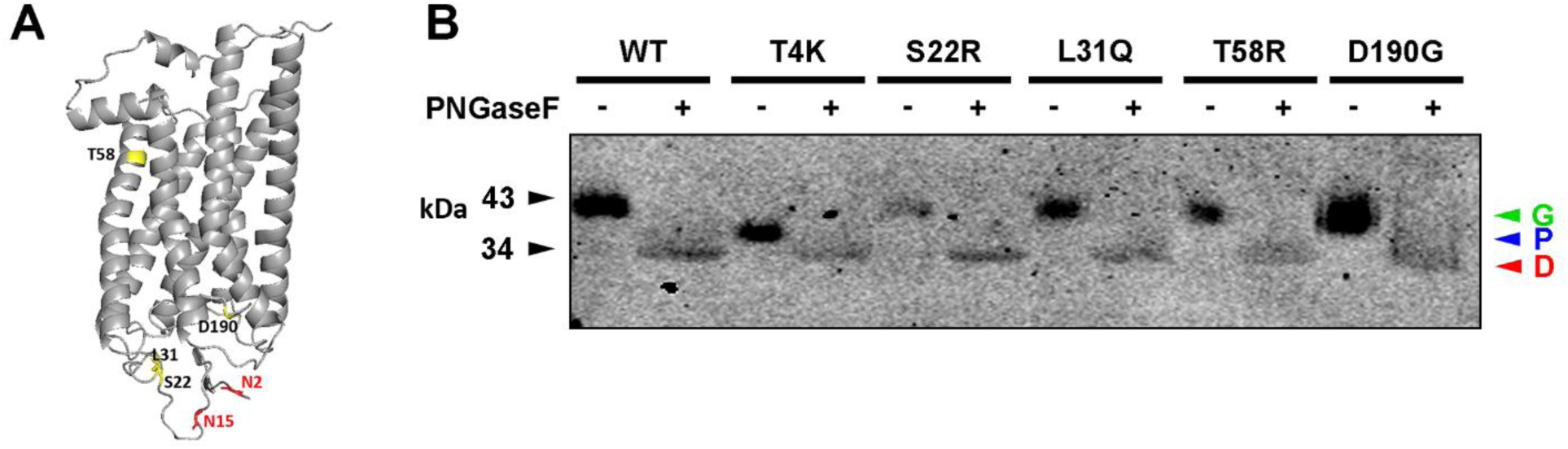
Class B1 RP-associated *RHO* mutations that do not alter consensus glycosylation sites do not affect RHO glycosylation. *A)* Location of RHO residues mutated (yellow) in relation to glycosylated residues (red). *B)* Western blot of solubilized transgenic *X. laevis* eyes expressing different human class B1 RP-associated RHO mutants with and without PNGaseF treatment. Western blot probed with mAb 1D4 (anti-mammalian RHO). “G” – glycosylated, “P” – partially glycosylated, “D” – deglycosylated.

### Dark rearing and preventing chromophore binding in *X. laevis* mitigate RD caused by ROS-localizing class B1 RHO mutants, but not RD caused by partially-ER retained RHO mutants

In animal models, dark rearing mitigates RD caused by class B1 mutations (specifically T4K, N15S, T17M, and P23H) but not other mutation classes, supporting a role for light as the environmental influence in class B1 RP.^18,41,43^ We examined the effects of light on two ROS-localizing *RHO* variants (L31Q or T58R) and two partially ER-retained variants (S22R or D190G). We raised transgenic *X. laevis* in constant dark or cyclic light and sacrificed them at 14 dpf. Dark rearing significantly reduced RD in animals expressing L31Q and T58R (ANCOVA: L31Q, p=0.01; T58R, p=0.0037) (Fig. 5A-D), but did not alter RD in animals expressing S22R or D190G (ANCOVA: S22R, p=0.4016; D190G, p=0.9569) (Fig. 5E-H).

**Figure 5.**
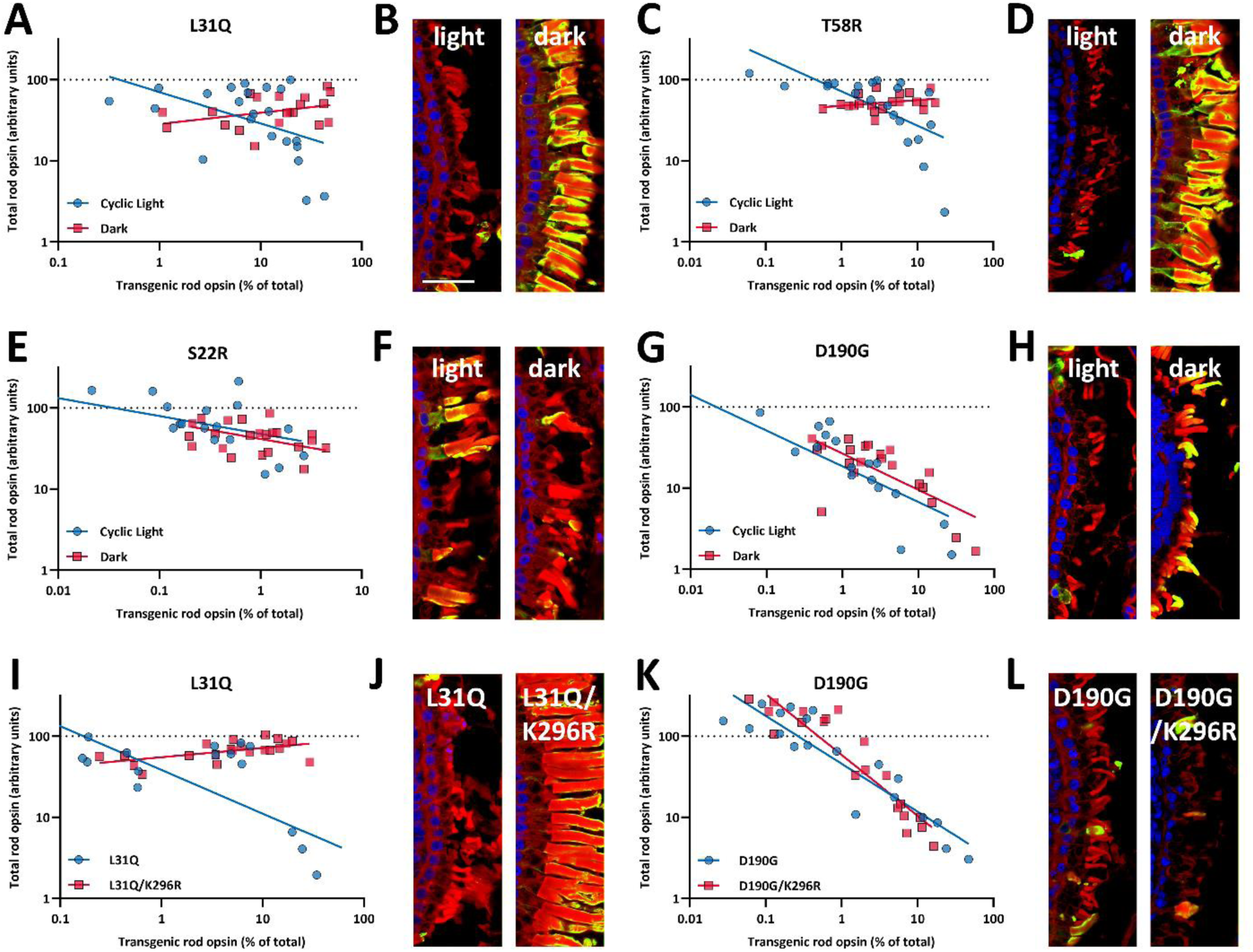
Effects of dark rearing and chromophore binding on degeneration in ROS-localizing and partially ER-retained class B1 variants. *A, C, E, G, I, K)* Dot blot graphs comparing total rod opsin expression versus percentage of transgenic rod opsin expression for ROS-localizing variants (L31Q, *A, I*; T58R, *C*) and partially RIS-localizing variants (S22R, *E*; D190G, *G, K*) examining either the effect of lighting condition (dark vs cyclic light) (*A-H*) or presence of bound chromophore (*I-L*) (normalized to WT standards, n = 18-22 animals). Solid line indicates linear regression of log transformed data. *B, D, F, H, J, L)* Confocal microscopy of representative retinas of 14 dpf transgenic *X. laevis* from the same experiment. Green – mammalian RHO (2B2), red – WGA, blue – Hoechst. Bar = 25 μm.

To determine whether it was necessary for these RHO variants to bind chromophore to cause light-exacerbated RD, as is the case for T4K and T17M but not P23H, we introduced a second mutation, K296R, that prevents chromophore binding but does not cause RD.^18,26,41,44^ We found that L31Q/K296R caused less degeneration than L31Q (ANCOVA: p=0.0003) (Fig. 5I-J). However, the toxicity of D190G/K296R was unaltered relative to D190G (ANCOVA: p=0.1133) (Fig. 5K-L).

### D190N *RHO* does not cause degeneration in the absence of light

We investigated the effects of light on the RD caused by D190N RHO, as it also lacked any significant mislocalization phenotype (Fig. 3A).^2^ This mutation was previously identified as both class A^45^ and class B1.^46^ Interestingly, dark-reared D190N animals had reduced RD relative to cyclic light-reared animals (*t-*test: p=0.0421) (Fig. 6A-B). Using our colocalization paradigm, we found no differences in D190N ER retention when comparing animals raised in cyclic light versus constant dark (*t*-test: p=0.4634) (Fig. 6C-D).

**Figure 6.**
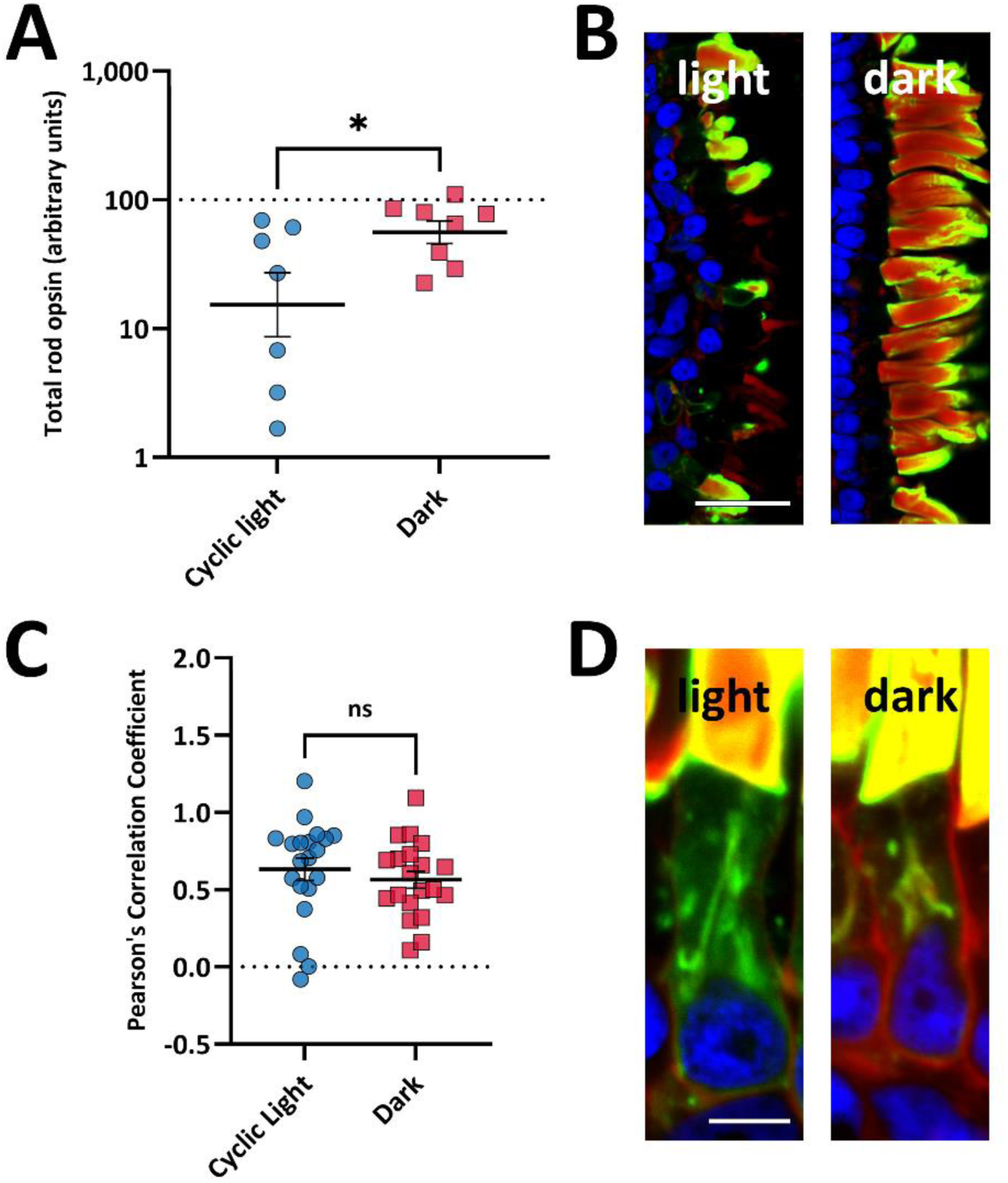
Effects of dark rearing on RD and trafficking of D190N RHO. *A)* Dot blot data comparing total rod opsin expression for animals expressing D190N *RHO* that were reared in cyclic light versus constant dark (normalized to WT standards, n = 7-8 animals). Error bars = SEM. *B)* Confocal microscopy of representative retinas of 14 dpf transgenic *X. laevis* raised in either cyclic light or constant dark. Green – mammalian RHO (2B2), red – WGA, blue – Hoechst. Bar = 25 μm. *C)* PCC values comparing trafficking of transgenic RHO in D190N *RHO*-expressing animals raised in cyclic light versus constant dark. N = 20 photoreceptors from two to three animals. Error bars = SEM. *D)* Confocal microscopy of representative retinas of transgenic *X. laevis* raised in either cyclic light or constant dark (green – mammalian RHO (2B2), red – WGA, blue – Hoechst). Bar = 5 μm.

### S176F RHO is retained in the ER and undergoes proteolytic cleavage

Because the localization of S176F closely resembled the class B1 mutant P23H, we investigated whether changes in light exposure altered its localization or toxicity. Using confocal microscopy, we confirmed RD in both cyclic light and dark-reared conditions (Fig. 7A). S176F was retained in RIS when labelled with an N-terminal antibody, where it overlapped with the ER marker calnexin and not with the Golgi marker WGA (Fig. 7B). To further verify RIS localization, we immunolabelled sections using anti-mammalian RHO antibodies directed at N- or C-terminal epitopes. N-terminal labeled S176F primarily mislocalized to RIS, with small quantities trafficked to the ROS that formed banded patterns, similar to previous results with P23H (Fig. 7C).^12,23^ However, C-terminally labeled S176F was primarily localized to ROS with some RIS retention, indicating that a fragment lacking the N-terminus trafficked to the ROS (Fig. 7D). To confirm this, we probed western blots of solubilized eye samples for S176F using both N- and C-terminal antibodies. Both antibodies labeled doublets at 18 kDa, indicating cleavage close to F176, near the center of the rod opsin sequence (Fig. 7E-F). The C-terminal antibody also labeled higher-order multimers or aggregates, which were observed in both lighting conditions (Fig. 7E-F).

**Figure 7.**
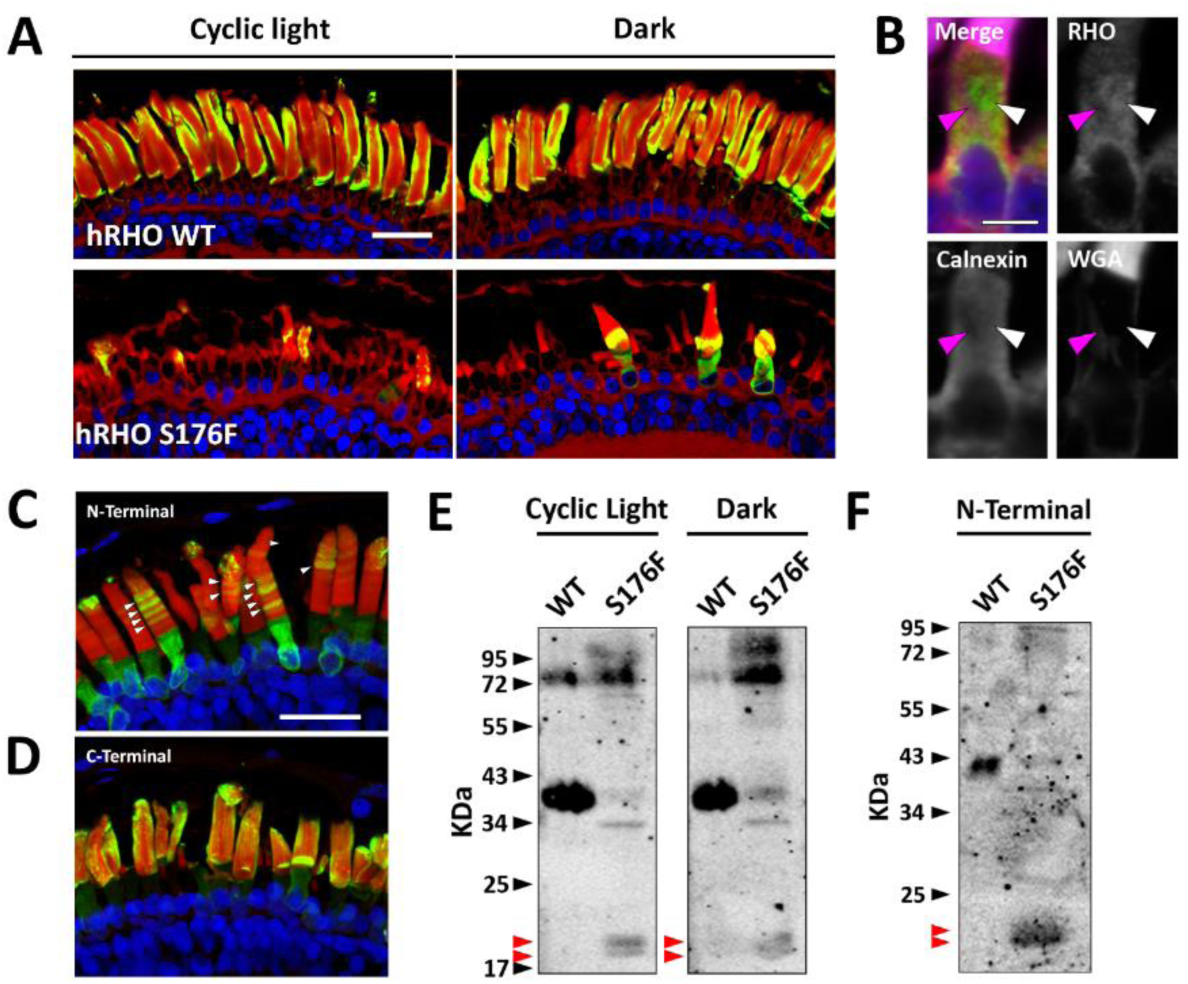
S176F RHO causes degeneration, undergoes proteolytic cleavage, and S176F fragments localize to different parts of the rod photoreceptors. *A)* Confocal images from cryosections of eyes from animals raised in either cyclic light or constant dark conditions. Red – WGA, green – mAb 2B2 (mammalian RHO), blue – Hoechst. Bar = 25 μm. *B)* High magnification image of a RIS in a primary S176F RHO transgenic animal. White arrows denote the presence of calnexin and transgenic RHO without WGA signal, while magenta arrows denote the presence of WGA without calnexin labelling and minimal RHO labelling. Magenta – WGA, green – anti-mammalian RHO (mAb 2B2), red – anti-calnexin, blue – Hoechst. Bar = 5 μm. *C-D)* Maximum projection images of two 12µm cryosections from the same F_0_ transgenic *X. laevis* S176F RHO eye labelled with either mAb 2B2 (N-terminal) (*C*) or mAb 1D4 (C-terminal) (*D*). White arrows denote bands of transgenic RHO in ROS. Red –WGA, green – anti-mammalian RHO (mAb 2B2 or mAb 1D4), blue – Hoechst. Bar = 25μm. *E-F)* Western blots of human WT and S176F RHO solubilized eye samples from animals raised cyclic light or constant dark. *E* – 1D4 (C-terminal), *F* – A5-3-12 (N-terminal). Red arrows denote cleaved fragments of S176F RHO.

## Discussion

We modelled RP caused by 26 *RHO* variants in transgenic *X. laevis* and conclusively observed RD in 19 models. Most mutants trafficked to the ROS similar to WT, while some displayed partial ER retention with ROS trafficking. Only S176F resembled the well-characterized P23H mutation, with observably low amounts of full length mutant RHO in the ROS, dramatic ER retention, and proteolytic cleavage. Class B1 mutants without localization defects caused photoactivation-dependent RD, while two partially ER-retained mutants did not. No additional class B1 *RHO* mutations investigated caused glycosylation deficiencies.

Most RP-associated *RHO* mutations we examined caused RD, including those causing photoactivation-induced toxicity (e.g. T4K), ER stress-related cell death (e.g. P23H), or plasma membrane mislocalization (e.g. A346P). However, none of the mutations at R135 caused RD, suggesting the associated mechanism is absent or non-pathogenic in *X. laevis*. A previous report suggests that R135 mutants have a defect in endocytosis, although a role for RHO endocytosis in vertebrate photoreceptors is not well established.^33^ Nevertheless, our results support a distinct mechanism for RD caused by these mutations. Differences in arrestin binding or endocytic responses in *X. laevis* may explain this paradox, warranting comparison in additional species.^47–49^ We also found no degeneration associated with F45L RHO; however, this mutation has been discredited as a cause of RP via population and patient data.^30,50–52^ Therefore, our system may prove useful in analyzing variants of uncertain significance, as *X. laevis* models can be rapidly generated for individual cases, with the caveat that some RD mechanisms may not be modeled.

Large photoreceptors and antibodies that preferentially label mammalian RHO enable detailed localization studies in *X. laevis*. Among 12 class B1 mutants, five displayed altered localization, indicating potentially novel pathogenic mechanisms distinct from ROS-localized mutants such as T4K and RIS-retained P23H. Notably, the pronounced mislocalization of P23H was mirrored only by the class A mutation S176F; statistically, these two variants constitute a homogeneous subset, suggesting both a distinct quality control mechanism, and the possibility that P23H should not be considered a “typical” RP mutation.

Dark rearing mitigated RD caused by ROS-localizing class B1 mutants (T4K, N15S, T17M, P23H, L31Q, and T58R), but not partially ER-retained class B1 mutants (S22R and D190G).^13^ Although lack of light-induced effects suggests that RD mechanisms involved in this group may be different between RP patients and our models, another possibility is that the two environmental conditions we compared were not suitable for resolving differences; notably, the degeneration induced by T4K is highly sensitive to the lighting paradigm, including intensity and cycle frequency;^25^ a different lighting paradigm may exist that would selectively exacerbate RD caused by S22R and/or D190G. Critically, RD caused by these mutants may not respond to T4K- or P23H-oriented therapeutic approaches due to mechanistic differences. In particular, D190G pathogenicity was not altered by preventing chromophore binding, suggesting that there is no “pharmacological chaperone” effect for these mutants.

Previously examined ROS-localizing class B1 mutations (T4K, T17M, and N15S) uniformly interfere with glycosylation. However, here we identified several examples of ROS-localizing variants that cause RD without altering glycosylation, including mutants that, based on reduced toxicity in dark reared animals and when chromophore binding is prevented, appear to share a similar cell death mechanism (e.g. L31Q). Thus, glycosylation defects may not be a prerequisite for this mechanism.^26^

In our model, D190N caused light-dependent RD, while patient data is conflicting on whether this mutation is associated with class A or class B1 RP. Unlike D190G, D190N shows no ER mislocalization, suggesting the light-induced RD caused by D190N is a ROS-originating phenotype similar to photoactivation-induced toxicity. Trafficking of D190N does not change between light and dark conditions, further signifying that misfolded transgenic RHO is not responsible for the sector phenotype. We predict that the correct clinical classification for D190N is class B1 RP. Several mutations have similarly been previously assigned to multiple clinical subclasses with T17M and E181K being classified as both class A and class B1 in different studies.^53^ This may reflect differences in the timing of phenotype reporting, as sector RP can progress to widespread retinal involvement over time, or different exposure to environmental influences (e.g. light) between patients.^54,55^ In the absence of sufficient patient data, animal models may assist in predicting the clinical class of *RHO* variants, and could help to inform physicians and researchers regarding treatment options. However, ultimately the clinical classification of mutations is based on human phenotypes, not animal models.

The class A-associated mutation S176F caused severe mislocalization and post-translational cleavage, similar to P23H, forming a unique group despite different clinical classifications. This suggests that while the cleavage and mislocalization occurs with both mutations, the severity of the molecular consequences such as protein misfolding (and its potential for prevention by chromophore binding) may determine the clinical phenotype. When examining the localization of S176F, it was apparent that the N-terminal fragments undergo quality control mechanisms and were retained in the ER, while the C-terminal fragments were trafficked to the ROS due to the intact ROS targeting motif and lack of the N-linked glycosylation that contributes to ER retention and quality control.^34,56^ Furthermore, the banding patterns of the N-terminal fragments suggests that this trafficking is moderated by a change that is repeated in regular intervals – likely the cyclical lighting and resulting pharmacological chaperone effect associated with higher chromophore availability in the dark.^12^ However, unlike P23H, this effect is insufficient to prevent RD.

Overall, our results emphasize the mechanistic heterogeneity of RD caused by RHO mutations, and the consequent need for a genotype-first approach in RP treatment. Therapeutic approaches that prevent RP caused by P23H may not be applicable to most other mutations. Differences between the T4K-like and P23H-like groups such as the remarkably different effects of vitamin A deprivation on RD suggest that treatments that have a positive effect on one group may have a negative effect on another.^2,26^ Clinical trials should stratify patients by mutation to optimize outcomes and identify mutation-specific therapies.

## Acknowledgements

We thank RS Molday for the gift of mAb 2B2 and WC Smith for mAbs B6-30N and A5-3-12.

---

**Supplementary Table S1.** AdRP-associated *RHO* mutations investigated in this study and previous studies using transgenic *X. laevis*. The table lists the mutation, residue domain (TM = transmembrane, EL = extracellular loop, CL = cytoplasmic loop), classification systems, proposed pathogenic mechanism, previously published animal models, *Xenopus* localization data, and evidence of light-induced degeneration in animal models. - indicates no available data. * based on Cideciyan et al.^10^. † based on Sung et al.^57^. ‡ based on Athanasiou et al.^30^. For localization, data is based on our colocalization assay (Fig. 3). ER+C – endoplasmic reticulum and cleavage, OS – outer segment, PER – partial ER retention, PM – plasma membrane.

| Mutation | Domain | Classification Systems |  |  | Animal Models | *Xenopus laevis*<br>Rod Localization | Light-Induced<br>Degeneration |
| --- | --- | --- | --- | --- | --- | --- | --- |
|  |  | Clinical* | Cell<br>Culture† | Pathogenic<br>Mechanism‡ |  |  |  |
| T4K | N-Terminus | B1 | II | 4 | *Xenopus*<sup>13</sup> | OS | Y<sup>13</sup> |
| N15S | N-Terminus | B1 | - | 2 | *Xenopus*<sup>13</sup> | OS | Y<sup>13</sup> |
| S22R<sup>58</sup> | N-Terminus | B1 | - | - | n/a | PER | N |
| T17M | N-Terminus | B1 | IIa | 4 | *Mouse*<sup>59</sup>,<br>*Xenopus*<sup>13</sup> | OS | Y<sup>13</sup> |
| P23H | N-Terminus | B1 | IIa | 2 | *Mouse*<sup>23,60,61</sup>,<br>*Pig*<sup>62</sup>, *Rat*<sup>63</sup>,<br>*Xenopus*<sup>18</sup> | ER+C | Y<sup>12</sup> |
| L31Q<sup>64</sup> | N-Terminus | B1 | II | - | n/a | OS | Y |
| M39R<sup>65</sup> | N-Terminus | B1 | I | 4 | n/a | OS | - |
| F45L<sup>28</sup> | N-Terminus | A | I | 7 | *Mouse*<sup>51</sup> | - | - |
| G51V<sup>66</sup> | TM1 | - | I | 2 | n/a | - | - |
| N55K<sup>67</sup> | TM1 | B1 | I | 4 | n/a | PER | - |
| T58R<sup>66</sup> | TM1 | B1 | IIb | 2 | n/a | OS | Y |
| L79P<sup>68</sup> | TM2 | - | - | - | n/a | PER | - |
| G89D<sup>66</sup> | TM2 | A | IIb | 2 | n/a | PER | - |
| L95P<sup>69</sup> | TM2 | B1 | - | 2 | n/a | OS | - |
| G101V<sup>70</sup> | EL1 | - | - | 2 | n/a | - | - |
| G106W<sup>28</sup> | EL1 | B1 | IIb | 2 | n/a | - | - |
| W126L<sup>71</sup> | TM3 | B1 | - | - | n/a | OS | - |
| R135G<sup>72</sup> | CL2 | A | IIb | 3 | n/a | OS | - |
| R135L<sup>73</sup> | CL2 | A | IIb | 3 | *Mouse*<sup>74</sup> | - | - |
| R135W<sup>28</sup> | CL2 | A | IIb | 3 | n/a | - | - |
| S176F<sup>75</sup> | EL2 | A | - | - | n/a | ER+C | - |
| P180A<sup>76</sup> | EL2 | B1 | - | - | n/a | - | - |
| P180T<sup>70</sup> | EL2 | - | - | - | n/a | OS | - |
| E181K<sup>66</sup> | EL2 | A | IIa | 2 | n/a | PER | - |
| D190G<sup>66</sup> | EL2 | B1 | IIa | 2 | n/a | PER | N |
| D190N<sup>28,77</sup> | EL2 | A / B1 | IIa | 2 | *Mouse*<sup>78</sup> | OS | Y |
| M207K<sup>79</sup> | TM5 | B1 | - | - | n/a | OS | - |
| P267L<sup>80</sup> | TM6 | B1 | IIa | 2 | n/a | PER | - |
| N310K<sup>69</sup> | CL4 | A | - | - | n/a | OS | - |
| Q344ter | C-Terminus | A | I | 1 | *Mouse*<sup>81</sup>,<br>*Xenopus*<sup>82</sup> | PM<sup>82</sup> | N<sup>43</sup> |
| A346P<sup>71</sup> | C-Terminus | A | - | 1 | n/a | PM | - |

**Supplementary Table S2.** Statistical results from comparisons of colocalization coefficients between RP-associated RHO mutants and human WT RHO. P-values from comparing PCC localization data from RP-associated mutants to either WT or P23H RHO using Dunnett’s multiple comparisons test. Data presented in Fig. 3.

| *RHO Variant* | *vs. WT* |  | *vs. P23H* |  |
| --- | --- | --- | --- | --- |
| WT | - |  | <0.0001 | **** |
| ***Class B1 RP*** |  |  |  |  |
| T4K | 0.8659 | ns | <0.0001 | **** |
| S22R | <0.0001 | **** | 0.0008 | *** |
| P23H | <0.0001 | **** | - |  |
| L31Q | 0.9894 | ns | <0.0001 | **** |
| M39R | 0.3849 | ns | <0.0001 | **** |
| N55K | 0.0141 | * | <0.0001 | **** |
| T58R | 0.9998 | ns | <0.0001 | **** |
| L95P | 0.6386 | ns | <0.0001 | **** |
| W126L | >0.9999 | ns | <0.0001 | **** |
| D190G | 0.0004 | *** | <0.0001 | **** |
| M207K | 0.3480 | ns | <0.0001 | **** |
| P267L | <0.0001 | **** | <0.0001 | **** |
| ***Class A RP*** |  |  |  |  |
| G89D | <0.0001 | **** | <0.0001 | **** |
| S176F | <0.0001 | **** | 0.6785 | ns |
| R135G | 0.3609 | ns | <0.0001 | **** |
| E181K | 0.0003 | *** | <0.0001 | **** |
| D190N | >0.9999 | ns | <0.0001 | **** |
| N310K | >0.9999 | ns | <0.0001 | **** |
| A346P | 0.9998 | ns | <0.0001 | **** |
| ***Other/Unknown RP*** |  |  |  |  |
| L79P | <0.0001 | **** | <0.0001 | **** |
| P180T | 0.9997 | ns | <0.0001 | **** |

